# Low-degree decompositions of flux vectors in faces of the flux cone

**DOI:** 10.1101/2024.05.22.595286

**Authors:** Frederik Wieder, Alexander Bockmayr

## Abstract

Decomposing a flux vector into elementary flux modes (EFMs) is a well-known task in the constraint-based analysis of metabolic networks.

Using geometric insights into the facial structure of the flux cone, we develop a new method for decomposition. To illustrate our approach, we consider a variety of genome-scale metabolic networks from the BiGG database. We decompose the optimal solutions to flux balance analysis problems into EFMs and compare different possible decompositions. Taking into account their degree allows understanding the interplay of the different EFMs that can participate in the decomposition of a given flux vector and highlights the particular importance of low-degree decompositions.

The new method can be applied to genome-scale metabolic networks, where the whole set of EFMs is often too large to be computed in practice. The special properties of low-degree decompositions make them an interesting subject for future biological studies.

## 1 Introduction

*Elementary flux modes* (EFMs) are an important concept in the constraint-based analysis of metabolic networks, see e.g. (Schuster and Hilgetag, 1994; Stelling et al, 2002; Schuster et al, 2002; Zanghellini et al, 2013). From the mathematical point of view, the EFMs provide an inner description of the steady-state flux cone of a metabolic network, which consists of a finite set of vectors such that any flux vector can be represented as a non-negative linear combination of EFMs (Gagneur and Klamt, 2004; Wagner and Urbanczik, 2005; Larhlimi and Bockmayr, 2008; Jevremovic and Boley, 2013; Röhl and Bockmayr, 2019). Even for small networks the number of elementary flux modes may be very large.

EFMs may be considered as minimal functional units of a metabolic network. Therefore, *decomposing a flux vector* into a non-negative linear combination of EFMs is an important task in metabolic network analysis see e.g. (Poolman et al, 2004; Schwartz and Kanehisa, 2005; Chan and Ji, 2011; Jungers et al, 2011; Rügen et al, 2012; Kelk et al, 2012; Chan et al, 2014; Maarleveld et al, 2015; Oddsdóttir et al, 2015; Gerstl et al, 2016; Chen et al, 2023). Particularly the flux distributions that optimize the flux through some given reaction, e.g. biomass, have been studied extensively as target vectors for the decomposition into EFMs. These optimizing vectors are typically obtained by performing a flux balance analysis (FBA), which is a linear optimization problem optimizing an objective function under a given set of linear constraints (Orth et al, 2010).

In a recent paper (Wieder et al, 2023), we studied the facial structure of the steady-state flux cone of a metabolic network and introduced the *degree* of a flux vector or EFM as the dimension of the inclusionwise minimal face of the flux cone containing this vector. The intuitive idea behind this concept is that the smaller the degree of an EFM, the more elementary it is. More formally, we showed that any flux vector of degree *d* > 0 can be decomposed into at most *d* EFMs of degree at most *t* + 1, where *t* is the dimension of the lineality space of the flux cone. In particular, any EFM of degree *d* ≥ *t* + 2 can be decomposed into at least two and at most *d* EFMs of degree strictly smaller than *d*. We also showed that any decomposition of a flux vector *v*\* only uses vectors that are contained in the face of the flux cone that is defined by *v*\*. In this paper, we use these mathematical results to develop a new method for decomposing flux vectors in genome-scale metabolic networks. We focus on decomposing optimal solutions *v*\* of flux balance analysis (FBA) problems. After computing *v*\*, we first determine the minimal face *F*\* of the flux cone *C* to which *v*\* belongs. Next we enumerate the EFMs in the face *F*\*, which typically has a much smaller dimension than the full flux cone. This way we are able to determine all the EFMs that may participate in a decomposition of *v*\*. Then we apply a mixed-integer linear programming (MILP) approach to compute different decompositions of *v*\* using these EFMs. Taking into account their degree allows understanding the interplay of the different EFMs that can participate in the decomposition of a given flux vector and highlights the particular importance of low-degree decompositions.

We illustrate our method on various genome-scale metabolic networks from the BiGG database (Norsigian et al, 2019a). For all these networks (with the exception of e coli core), we are not able to compute the full set of EFMs, while the EFMs in the face *F*\* can be obtained easily. Their number is typically rather small, which makes it possible to explore the different possible EFM decompositions with MILP. While there exist often many different shortest EFM decompositions of a given target vector *v*\*, it turns out that *low-degree decompositions*, using only EFMs of degree at most *t* + 1, in many cases are unique and provide the building blocks for the other decompositions.

## 2 Background

A *metabolic network* 𝒩 = (ℳ, ℛ, *S*, Irr) is given by a set ℳ of (internal) *metabolites*, a set 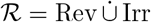 of *reversible* and *irreversible reactions*, and a *stoichiometric matrix S* ∈ ℝ^*m×n*^, where *m* = |ℳ| and *n* = |ℛ|. The network can be seen as a weighted hypergraph with the generalized incidence matrix *S*, where the metabolites are represented by nodes and the reactions by hyperarcs. A positive entry *S*_*i,j*_ > 0 in the stoichiometric matrix *S* indicates that reaction *j* produces metabolite *i*. If *S*_*i,j*_ < 0, metabolite *i* is consumed in reaction *j*.

We assume that the network is in *steady-state*, i.e., for each internal metabolite the rate of production is equal to the rate of consumption. In matrix notation, the steady-state constraints can be written as *Sv* = 0, where *v* ∈ ℝ^*n*^ denotes a *flux vector*. By adding the thermodynamic irreversibility constraints *v*_Irr_ ≥ 0, where *v*_Irr_ is the subvector of *v* whose components belong to Irr, we obtain the polyhedral cone

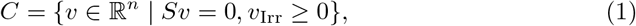

which is called the (steady-state) flux cone of 𝒩.

A flux vector *v* ∈ *C* is *reversible* if and only if both *v* and −*v* belong to *C*. The *lineality space L* of *C* is the set of all reversible flux vectors and can be described as

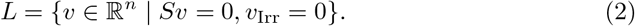

In metabolic network analysis we are particularly interested in minimal functional units of the network. A flux vector *e* ∈ *C \* {0} is called an *elementary flux mode* (EFM) if it has inclusionwise minimal support, i.e., if

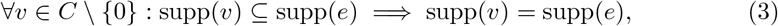

where the *support* of *v* ∈ ℝ^*n*^ is defined by supp(*v*) = {*i* ∈ ℛ | *v*_*i*_*≠* 0}. We say that a reaction *i* ∈ ℛ is *active* in *v* ∈ *C* if *i* ∈ supp(*v*). Any vector *v*\* ∈ *C* in the flux cone *C* of a metabolic network can be decomposed into a positive linear combination

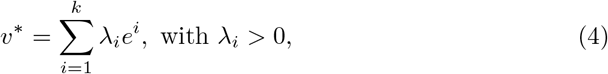

of elementary flux modes *e*^1^, …, *e*^*k*^ ∈ *C*, see e.g. Lemma 1 in (Schuster et al, 2002). By requiring *λ*_*i*_ > 0 we can define the *length* of the decomposition as *k* and a *shortest decomposition* as a decomposition of minimal length. Note that in most cases shortest decompositions are not unique (cf. Fig. 4).

For any *J ⊆* Irr, the set *F*_*J*_ = {*v* ∈ *C* | *v*_*j*_ = 0 for all *j* ∈ *J*} is called a *face* of the polyhedral cone *C*. Given *v*\* ∈ *C*, the inclusionwise minimal face of *C* containing *v* is *F*_*J*(*v*_*∗*_)_ with 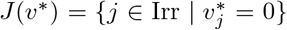. We call *F*\*.= *F*_*J*(*v*_*∗*_)_ *⊆ C* the *face defined by v*\*

It was shown in (Wieder et al, 2023) that any decomposition (4) of *v*\* ∈ *C* only includes vectors from the face *F*\* defined by *v*\*. More formally, if 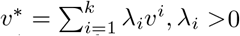 0, is a decomposition of a flux vector *v*\* ∈ *C* into flux vectors *v*^1^, …, *v*^*k*^ ∈ *C*, then *v*^1^, …, *v*^*k*^ ∈ *F*\*, where *F*\* is the face defined by *v*\*.

The *degree* deg(*v*) of a flux vector *v* ∈ *C* was introduced in (Wieder et al, 2023) as the dimension of the face *F*_*J*(*v*)_ defined by *v*. We have

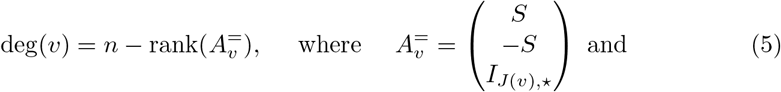

*I*_*J*(*v*),*⋆*_ is the submatrix of the (*n×n*) identity matrix *I*_*n*_ consisting of the rows belonging to *J* (*v*). The intuition underlying the concept of degree is that the smaller the degree, the more elementary the EFM or more generally the flux vector is.

It was shown in (Wieder et al, 2023) that any flux vector *v*\* ∈ *C* of degree *d* = deg(*v*\*) > 0 has a decomposition 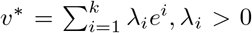, into at most *k* ≤ *d* EFMs *e*^1^, …, *e*^*k*^ of degree at most *t* + 1, where *t* = dim(*L*) is the dimension of the lineality space *L* of *C*. We call such a decomposition a *low-degree decomposition*. Note that the EFMs of degree *t* or *t* + 1 participating in a low-degree decomposition are elements of a *minimal set of elementary modes* (MEMo), i.e., a minimal set of EFMs generating the flux cone *C* (Röhl and Bockmayr, 2019). Therefore, EFMs of degree *t* and *t* + 1 can also be called *MEMo-EFMs*.

As a special case, we get that any EFM *e* ∈ *C* with deg(*e*) ≥ *t* + 2 has a decomposition into at least two and at most *d* EFMs of degree strictly smaller than *d*.

The following example and the results of Tab. 2 in Sect. 4 show that shortest decompositions of flux vectors or EFMs typically contain EFMs of degrees larger than *t* + 1, i.e., they are usually not low-degree decompositions.

### An illustrating example

The metabolic network in Fig. 1 has the set of metabolites ℳ = {A, B, …, G}, the set of reversible reactions Rev = {1, 3, 4, 5, 9, 10, 11, 12} and the set of irreversible reactions Irr = {2, 6, 7, 8}. The stoichiometric matrix is

**Fig. 1.**
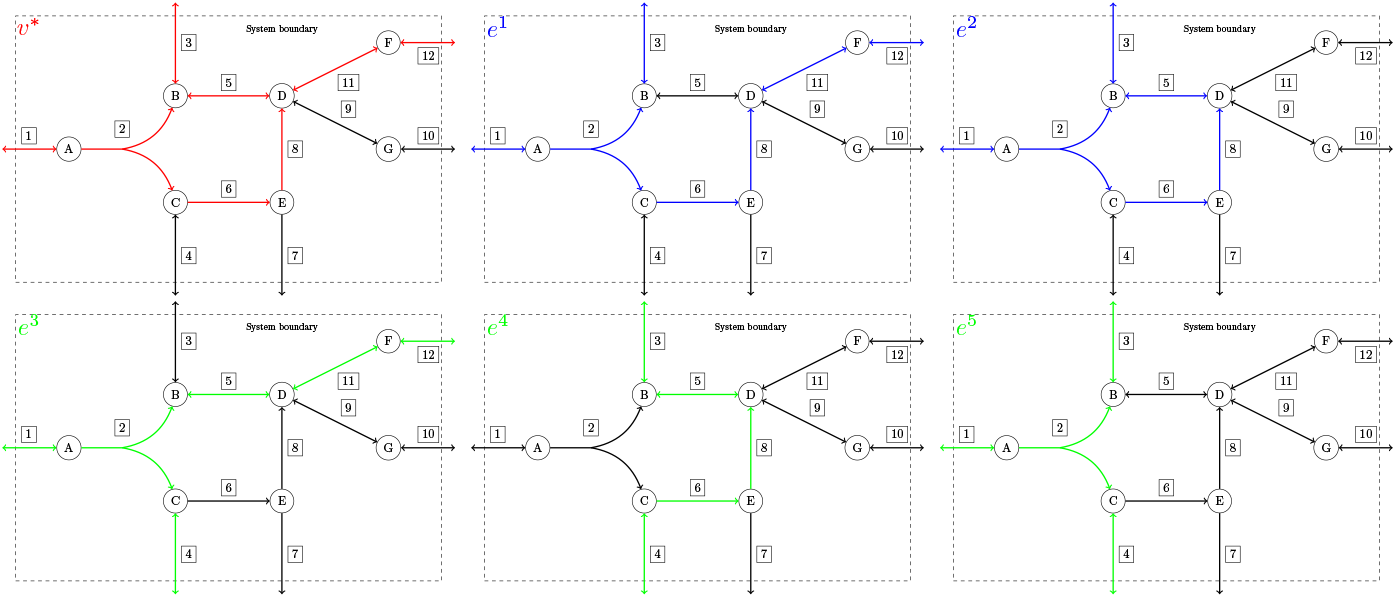
Vector *v*\* to decompose (red), EFMs *e*^1^, *e*^2^ of degree 4 (blue) and EFMs *e*^3^, *e*^4^, *e*^5^ of degree 3 (green) for a small example network.

**Fig. 2.**
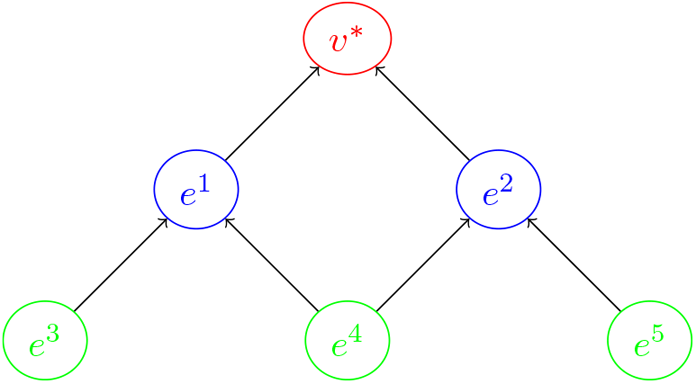
Decomposition of *v*\* into two EFMs of degree 4 (blue), which can be further decomposed into 3 MEMo-EFMs of degree 3 (green).

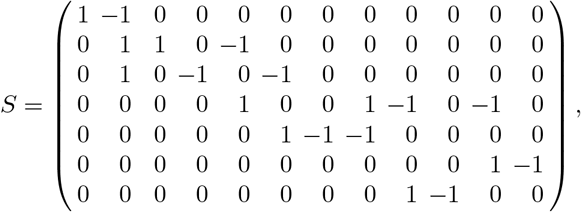

where we assume that reversible reactions are oriented from left to right and from top to bottom and only chose stoichiometric coefficients from -1,0 and 1.

The vector *v*\* (in red) is the one to be decomposed and has degree 4. It could be either a given steady-state flux vector or the optimal solution of some FBA problem (e.g. maximizing flux rates of reactions 3 and 12). In a first step, a shortest decomposition of *v*\* into EFMs is computed. The resulting EFMs *e*^1^ and *e*^2^ (in blue) have degree 4 and it holds *v*\* = *e*^1^ + *e*^2^. In the next steps, *e*^1^ and *e*^2^ are decomposed into EFMs of lower degree. The resulting EFMs *e*^3^, *e*^4^, *e*^5^ (in green) have degree 3 and cannot be further decomposed into EFMs of smaller degrees. It holds *e*^1^ = *e*^3^ + *e*^4^ and *e*^2^ = *e*^4^ + *e*^5^, and thus *v*\* = *e*^3^ + 2*e*^4^ + *e*^5^. The corresponding vectors are (the symbol ·^*T*^ denotes transposition)

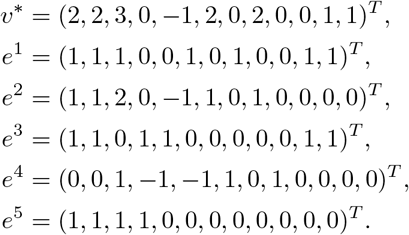

In this simple example, the weights *λ*_*i*_ in the above decompositions are all small integers. In general, this need not be the case. In particular, the weights will depend on the normalization or scaling of the EFMs, which are determined only up to multiplication by a positive number.

## 3 Methods

As we have seen in Sect. 2, for decomposing a flux vector *v*\* ∈ *C* into EFMs it suffices to consider those EFMs that are contained in the face *F∗* = *F*_*J*(*v*_*∗*_)_ *⊆ C* defined by *v*\*. The face *F*\* is again a polyhedral cone, which informally is the flux cone of the metabolic network 𝒩^*′*^ = (ℳ, ℛ *\ J* (*v*\*), *S*^*′*^, Irr^*′*^) obtained from 𝒩 = (ℳ, ℛ, *S*, Irr) by deleting the reactions in *J* (*v*\*).

By computing the EFMs of *N*^*′*^ using standard tools like *efmtool* (Terzer, 2009) or *EFMlrs* (Buchner and Zanghellini, 2021) (we used *efmtool*), and adding zero components *v*_*j*_ = 0 for the deleted reactions *j* ∈ *J* (*v*\*), we obtain exactly the EFMs of *N* contained in *F*\*. For a formal proof, we refer to Lemma 4 in (Marashi and Bockmayr, 2011).

Our workflow for decomposing the optimal solution of an FBA problem into EFMs is shown in Fig. 3. The main steps are as follows:

**Fig. 3.**
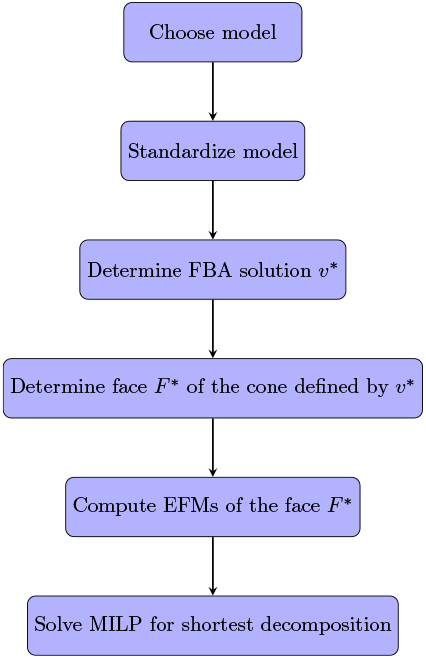
Workflow for determining decompositions of an FBA solution into EFMs

### Choose model

After downloading an SBML (Norsigian et al, 2019b) file from the BiGG database, the software package `cobrapy` (Ebrahim et al, 2013) offers the function `cobra.read sbml model()` to create a model from the file containing all metabolites, reactions, bounds and further information from the chosen metabolic network.

### Standardize model

In their standard (downloadable) form, some models in the BiGG database are not suitable for EFM computation with *efmtool*. This is due to the fact that sometimes irreversible uptake reactions are oriented as negative output reactions (i.e., an irreversible reaction has a negative flux, which is not in line with the requirements of *efmtool*). This can be fixed by changing the orientation of these reactions. We did this by replacing these reactions with correctly oriented copies.

Furthermore, we removed all reactions where the lower and upper bound are both equal to zero.

### Determine FBA solution v*

The standard objective is to maximize biomass production. By calling the function `cobra.optimize()`, a flux vector *v*\* achieving the optimal growth rate is computed. We did not change any objective function and just worked with the model exactly the way it can be downloaded (up to changing the orientation of some irreversible exchange reactions and removing reactions with lower and upper bound equal to 0, as mentioned above). If one is interested in decomposing some flux vector that is not obtained by solving an FBA problem, this step can be skipped and the given vector can be used as *v*\* instead of the FBA solution.

### Determine the face F* defined by v*

By calling the function `cobra.util.create stoichiometric matrix()`, the stoichiometric matrix of the flux cone *C* can be obtained. Adding unit row vectors to the stoichiometric matrix corresponding to the constraints *v*_*j*_ = 0, *j* ∈ *J* (*v*\*), together with a 0-1-vector containing information of the reversibility of reactions, leads to a description of the face *F*\* containing all candidate EFMs for decomposing the target vector *v*\*.

### Compute EFMs of the face F*

The modified stoichiometric matrix and the reversibility vector are used as input for EFM computation with *efmtool* (Terzer, 2009). The EFMs computed in this step represent all EFMs in the face *F*\* of the flux cone that contains the optimal FBA solution *v*\*. They form a subset of the set of all EFMs in the flux cone.

### Solve MILP for shortest decomposition

A shortest decomposition of a vector *v*\* given a set of candidate vectors (in our case EFMs in *F*\* or MEMo-EFMs in *F*\*) can be found by solving a mixed-integer linear program (MILP):

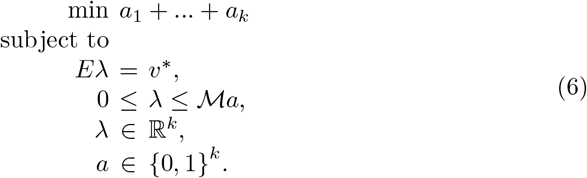

Here, *E* = (*e*^1^, …, *e*^*k*^) ∈ ℝ^*n×k*^ is the matrix containing the *k* candidate EFMs *e*^1^, …, *e*^*k*^ in the face *F*\* as columns. The vector *λ* ∈ ℝ^*k*^ contains the coefficients of the EFMs in the decomposition *Eλ* = *λ*_1_*e*^1^ + ⋯ + *λ*_*k*_*e*^*k*^ = *v*\*. The binary variables *a*_*j*_ indicate whether or not EFM *e*^*j*^ takes part in the decomposition, i.e., whether *λ*_*j*_ > 0 or *λ*_*j*_ = 0. By minimizing the sum *a*_1_ +⋯ +*a*_*k*_, we find a shortest decomposition using as few EFMs as possible. The bigM constant ℳ is an integer number that has to be chosen sufficiently large (we used 10^6^ for the calculations in Tab. 2). The constraint 0 ≤ *λ* ≤ ℳ*a* implies that *λ*_*j*_ = 0 if *a*_*j*_ = 0 and that *λ*_*j*_ is unbounded (from above) if *a*_*j*_ = 1 (more precisely, *λ*_*j*_ is bounded by ℳ, that is why choosing ℳ large enough is crucial). ^1^

In general, different decompositions can be computed with the set of EFMs contained in the face *F*\*, depending on their degree. We will illustrate this in Sect. 4 and Sect. 5 by presenting and discussing not only shortest decompositions considering all EFMs in *F*\*, but also shortest decompositions into MEMo-EFMs. For shortest decompositions into MEMo-EFMs, the set of candidates in the decomposition is further restricted to EFMs of degree *t* := dim(*L*) or *t* + 1.

## 4 Results

After having introduced our method in Sect. 3, we present now computational results for decomposing optimal solutions of FBA problems from the BiGG database (Norsigian et al, 2019a). All computations were performed on a Thinkpad T14s with an AMD Ryzen 7 PRO 4750U Processor and 32GB of RAM.

Tab. 1 shows for a large selection of genome-scale metabolic networks from the BiGG database the number of EFMs to be considered for decomposing the optimal FBA solution of the given model and the time needed to compute these EFMs. These running times range between a few seconds and a couple of minutes. This is remarkable because for all but one of these networks we are unable to compute the full set of EFMs on our machine, due to memory limitations. The only exception is e coli core, which has a total number of 100274 EFMs.

**Table 1.**
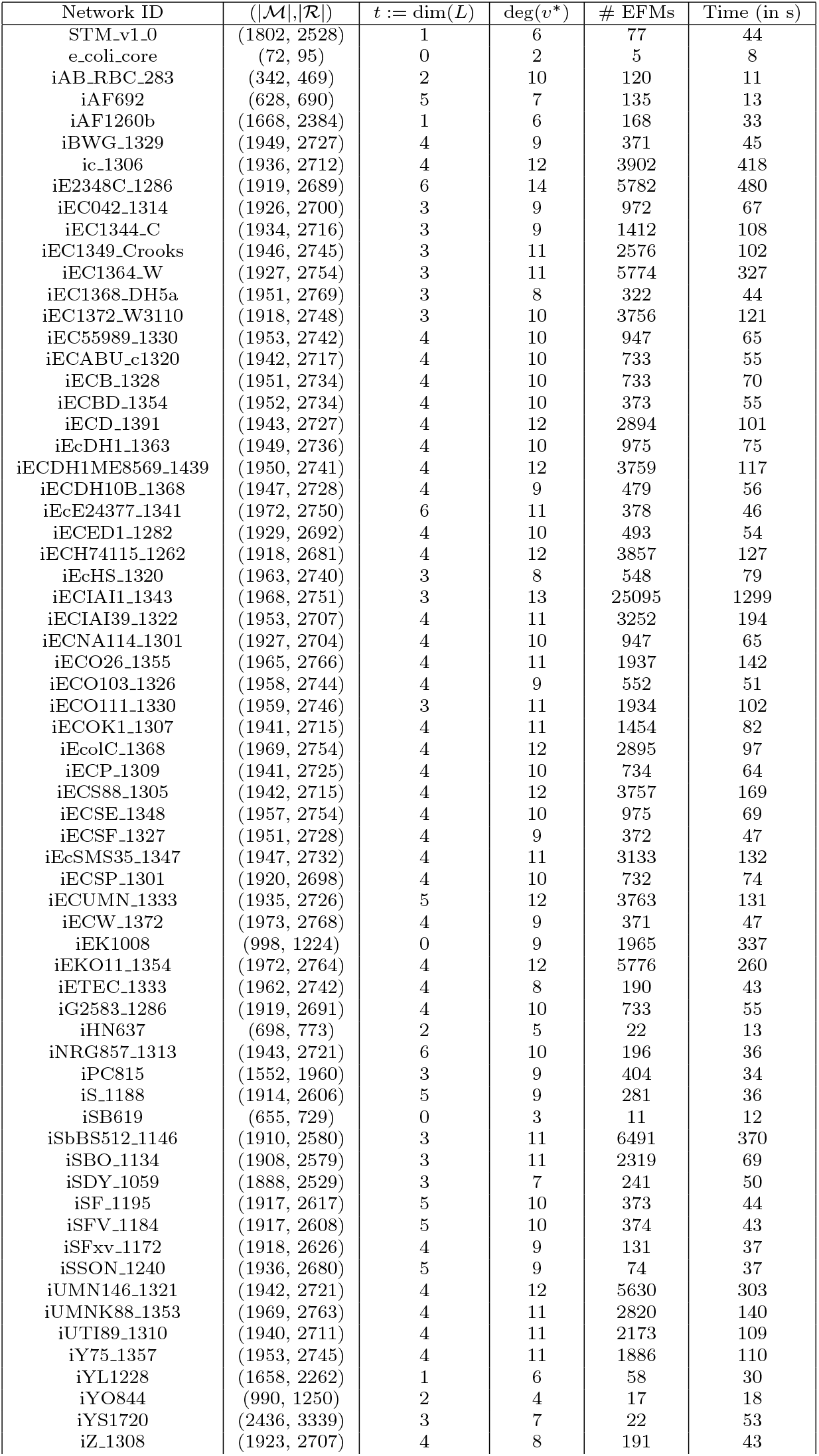
EFMs in the face *F*\* defined by the optimal solution *v*\* of FBA problems in the BiGG database. Network Id: Identifier of the network; (|ℳ|,|ℛ|): Number of metabolites and reactions; t: Dimension of the lineality space; deg(*v*\*): Degree of the optimal FBA solution *v*\*. # EFMs: Number of EFMs in the face *F*\* defined by *v*\*. Time: Running time (in seconds) to compute the EFMs in *F*\*.

The first column in Tab. 1 contains the identifier of the model in the BiGG database. The second column represents the size of the metabolic network (after our standardization) by stating the number of metabolites and reactions in the network. Column 3 gives the dimension *t* of the lineality space, which is relatively small for all the networks, ranging between 0 and 6 (if *t* = 0, the corresponding flux cone is pointed). The degree of the optimal FBA solution *v*\* provided by `cobrapy` (Ebrahim et al, 2013) is given in column 4. Again these degrees are relatively small and range between 2 and 12. Note that the dimension of the full flux cone is typically much higher. Due to the small degree of *v*\*, which equals the dimension of the face *F* ^*∗*^ defined by *v*\*, it becomes possible to compute all EFMs contained in this face. Their number is given in column 5. As explained in Sect. 2, there are no other EFMs needed to decompose *v*\*. Finally, column 6 contains the running time (in seconds) to compute all these EFMs, which is less than 10 minutes for all but 1 network (iECIAI1 1343 with 25095 EFMs).

Tab. 2 provides more detailed information about the EFMs and the shortest decompositions for a subset of the models in Tab. 1. The first four columns are the same as before. Column 5 lists the degrees of the EFMs in the shortest decomposition computed by the MILP (6), while column 6 lists the degrees of EFMs in the shortest decomposition into MEMo-EFMs. For example, in the first row of Tab. 2, the optimal solution to the FBA problem for STM v1 0 has degree 6 and it can be decomposed into two EFMs of degree 5 or 5 EFMs of degree 2. Finally, the last column contains a list summarizing information about the degrees of all EFMs in the face *F*\* defined by *v*\*. Looking again at the first row, the list [0,2,5,12,26,24,8] should be read in the following way: The face containing the optimal solution contains no EFMs of degree 0 (this immediately follows from the fact *t* := dim(*L*) = 1). There are two EFMs of degree 1, five EFMs of degree 2, 12 of degree 3, 26 of degree 4, 24 of degree 5 and 8 of degree 6.

**Table 2.** Decompositions of FBA solutions *v*\* for a selection of networks from the BiGG database. (|ℳ|,|ℛ|): Number of metabolites and reactions, t: Dimension of the lineality space, deg([EFMs]): Degrees of EFMs in shortest decomposition, deg([MEMo-EFMs]): Degrees of EFMs in shortest MEMo-EFM decomposition, EFM distribution in *F*\*: Number of EFMs with degree equal to their index in the list.

| Network ID | $(\vert \mathcal{M}\vert , \vert \mathcal{R}\vert )$ | t | $\deg(v^*)$ | $\deg(\text{EFMs})$ | $\deg(\text{MEMo-EFMs})$ | EFM distribution in $F^*$ |
| --- | --- | --- | --- | --- | --- | --- |
| STM_v1.0 | (1802, 2528) | 1 | 6 | [5,5] | [2,2,2,2,2] | [0, 2, 5, 12, 26, 24, 8] |
| e.coli_core | (72, 95) | 0 | 2 | [1,2] | [1,1] | [0, 2, 3] |
| iAB.RBC.283 | (342, 469) | 2 | 10 | [5,4,4,4,4,4,4] | [3,3,3,3,3,3,3,3] | [0, 0, 4, 18, 44, 54, 0, 0, 0, 0, 0] |
| iAF1260b | (1668, 2384) | 1 | 6 | [6,5] | [2,2,2,2,2] | [0, 2, 11, 32, 59, 49, 15] |
| iLJ478 | (570, 652) | 2 | 5 | [5] | [3,3,3] | [0, 0, 6, 15, 21, 7] |
| iSB619 | (655, 729) | 0 | 3 | [2,2] | [1,1,1] | [0, 3, 6, 2] |
| iSSON_1240 | (1936, 2680) | 5 | 9 | [9,8] | [6,6,6,6] | [0, 0, 0, 0, 0, 12, 8, 21, 24, 9] |

## 5 Discussion

We now further discuss the computational results from Sect. 4. Consider again the first row of Tab. 2. In total, there are only 7 MEMo-EFMs, and the shortest decomposition into MEMo-EFMs we computed uses all five EFMs of degree 2. Fig. 4 visualizes how the two EFMs of degree 5 in the shortest decomposition can be further decomposed into the five EFMs of degree 2 in the shortest MEMo-decomposition. In Fig. 4, the arrows pointing to an EFM start at all EFMs that are used in a decomposition of it. So the two arrows pointing from the two EFMs of degree 5 to the FBA solution of degree 6 indicate that the optimal solution of degree 6 can be decomposed into two EFMs of degree 5. Furthermore, each of the EFMs of degree 5 can be decomposed into one EFM of degree 4 and one of degree 2, and so on.

**Fig. 4.**
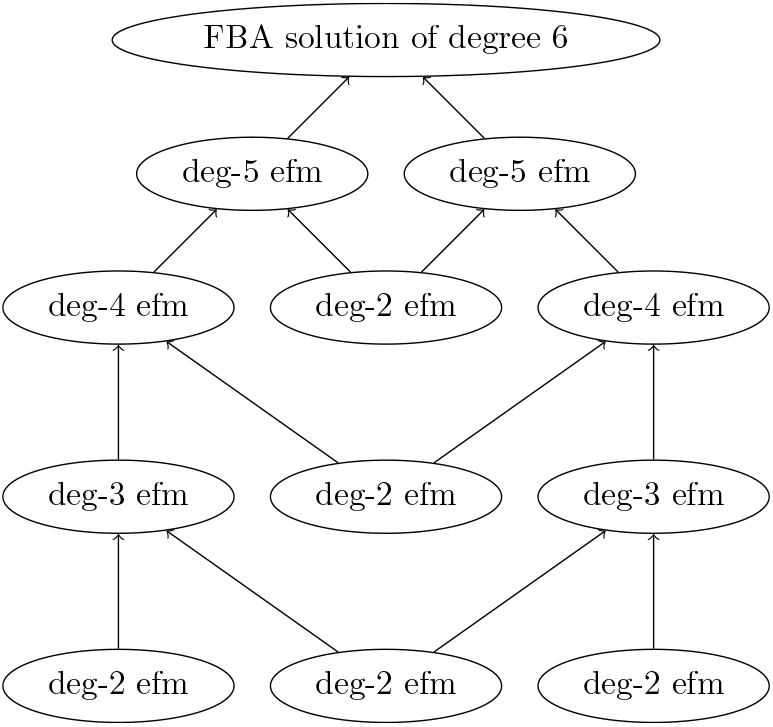
Decomposition of the optimal FBA solution *v*\* of STM v1 0 into MEMo-EFMs.

From Tab. 1 and Tab. 2, we can immediately see that, by determining the face *F*\* defined by the optimal solution *v*\*, we are able to find all candidate EFMs for a decomposition in metabolic networks where the enumeration of all EFMs is not possible in practice. In fact, we were only able to compute all 100274 EFMs of e coli core, while the computation of all EFMs for any of the other networks was not possible on our computer, due to memory limitations.

The advantage of determining shortest decompositions into MEMo-EFMs lies in the fact that these EFMs cannot be further decomposed into EFMs of smaller degrees. Thus, the MEMo-EFMs in the low-degree decompositions are the building blocks for the decompositions of higher degree.

Furthermore, the number of different possible shortest decompositions becomes much smaller when using MEMo-EFMs, as Tab. 3 shows. For all the networks in Tab. 3, the shortest decomposition into MEMo-EFMs turns out to be unique, while there exist multiple shortest decompositions into EFMs of higher degree (with the exception of iLJ478).

**Table 3.** Number of different shortest decompositions into EFMs and MEMo-EFMs.

| Network ID | # shortest decompositions | # shortest decompositions<br>into MEMo-EFMs |
| --- | --- | --- |
| STM_v1_0 | 5 | 1 |
| e.coli_core | 4 | 1 |
| iAB.RBC_283 | 12 | 1 |
| iAF1260b | 4 | 1 |
| iLJ478 | 1 | 1 |
| iSB619 | 2 | 1 |
| iSSON_1240 | 4 | 1 |

To determine all different shortest decompositions we can solve iteratively the MILP (6), adding each time a so-called *no-good cut* of the form

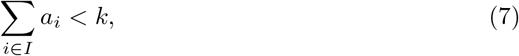

where 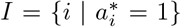 is the set of indices and *k* the length of the shortest decomposition (*λ*\*, *a*\*) found in the previous step. This allows enumerating all shortest decompositions, using a different set of EFMs in each step. When there are no more shortest decompositions of length *k*, the MILP will compute decompositions of higher length.

Although in Tab. 3 all shortest decompositions into MEMo-EFMs are unique, this need not always be the case, as the following example shows. Consider a slightly modified version of iAF 1260b, namely iAF 1260, again from the BiGG database. After performing an FBA, the resulting flux vector *v*\* has degree 7 and can be decomposed into two EFMs of the degrees 6 and 7. The 7-dimensional face defined by *v*\* contains 546 EFMs. Of those, only 2 are in the lineality space (have degree 1), and 16 have degree 2, resulting in a total of 18 MEMo-EFMs. Using these 18 MEMo-EFMs as candidates, there exist two different shortest decompositions of the FBA solution *v*\*, which both have length 6.

A visual representation why there can be multiple shortest decompositions is given in Fig. 5. Consider e coli core, where the degree of the optimal FBA solution *v*\* is 2 and only 5 EFMs are contained in the face *F*\* defined by *v*\*. The figure shows a representation (not to scale) of the face *F*\*. The optimal solution *v*\* can be decomposed into the EFM *e*^2^ and any one of the other four EFMs. A decomposition of *v*\* without *e*^2^ is not possible. Furthermore, the EFMs *e*^3^, *e*^4^ and *e*^5^ of degree 2 can be decomposed into the extreme rays of the cone, which are the EFMs *e*^1^ and *e*^2^ of degree 1. In summary, any vector in Fig. 5 that lies between two others can be decomposed into them.

**Fig. 5.**
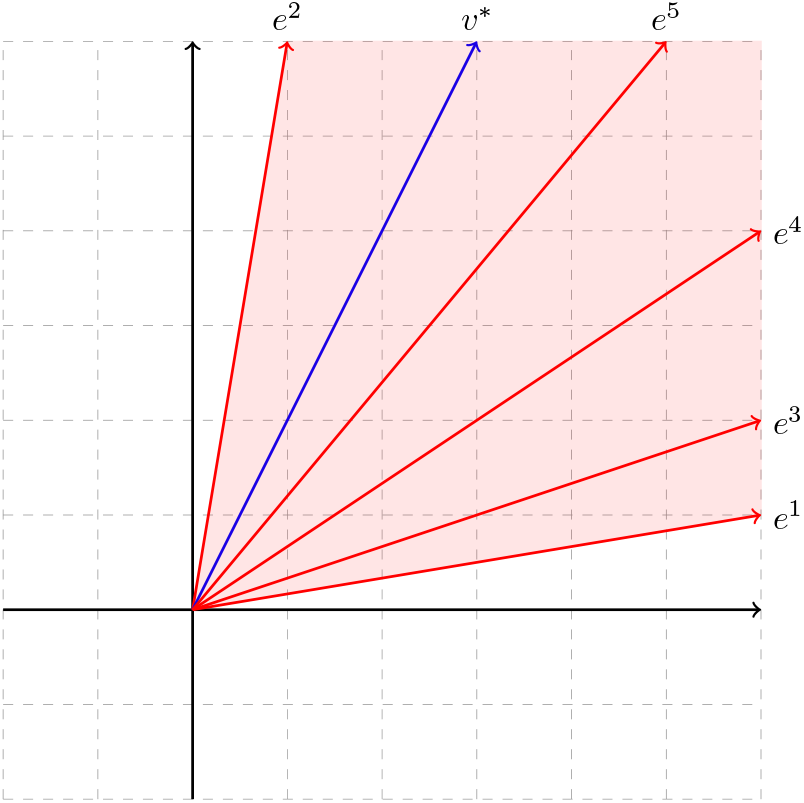
Visualization of the 2-dimensional face *F*\* defined by the optimal FBA solution *v*\* of e coli core and the 5 EFMs contained in it.

## 6 Conclusion

Based on geometric insights into the facial structure of the flux cone of a metabolic network, we developed a novel method for decomposing flux vectors into elementary flux modes (EFMs). While our approach is applicable to any given flux vector in the cone, we focused on common decomposition targets, namely solutions to FBA problems. By reducing the search space for EFM computation to the face of the flux cone defined by the target vector, the number of EFMs to be computed becomes much smaller. Therefore, it can also be applied to genome-scale metabolic networks where computing all EFMs is not possible in practice.

Taking into account the degree of the EFMs, we obtained additional insights into the structure of different decompositions and highlighted the special role of low-degree decompositions consisting of MEMo-EFMs, which cannot be further decomposed into EFMs of smaller degree and thus are the building blocks for the decompositions of higher degree.

We illustrated our approach on various genome-scale metabolic networks from the BiGG database. For all these networks, the EFMs in the face defined by the optimal FBA solution can be obtained easily and it is possible to enumerate different shortest decompositions either into MEMo-EFMs or EFMs of higher degree. In many cases, the low-degree decomposition in MEMo-EFMs turns out to be unique, although this need not be the case in general, Future research should address the biological implications of low-degree decompositions and the MEMo-EFMs on which they are based.

## Acknowledgement

The authors want to thank Guillermo Garcia Martínez for implementing the MILP.

## Footnotes

1 We used the “max” normalization option offered by *efmtool*, which scales the computed EFMs to have a largest absolute value of 1. Using the default “min” normalization option that scales the smallest absolute value to 1 leads to numerical issues when solving the MILP for some models.

